# Divergent profiles of fentanyl withdrawal and associated pain in mice and rats

**DOI:** 10.1101/2020.11.16.384818

**Authors:** Olivia Uddin, Carleigh Jenne, Megan E. Fox, Keiko Arakawa, Asaf Keller, Nathan Cramer

**Author notes:** The first two authors contributed equally to the work described in this manuscript. Address Correspondence to: Dr. Nathan Cramer, 20 Penn Street, Baltimore, MD, 21201.

## Abstract

Opioid abuse has devastating effects on patients, their families, and society. Withdrawal symptoms are severely unpleasant, prolonged, and frequently hinder recovery or lead to relapse. The sharp increase in abuse and overdoses arising from the illicit use of potent and rapidly-acting synthetic opioids, such as fentanyl, highlights the urgency of understanding the withdrawal mechanisms related to these drugs. Progress is impeded by inconsistent reports on opioid withdrawal in different preclinical models. Here, using rats and mice of both sexes, we quantified withdrawal behaviors during spontaneous and naloxone-precipitated withdrawal, following two weeks of intermittent fentanyl exposure. We found that both mice and rats lost weight during exposure and showed increased signs of distress during spontaneous and naloxone precipitated withdrawal. However, these species differed in their expression of withdrawal associated pain, a key contributor to relapse in humans. Spontaneous or ongoing pain was preferentially expressed in rats in both withdrawal conditions, while no change was observed in mice. In contrast, withdrawal associated thermal hyperalgesia was found only in mice. These data suggest that rats and mice diverge in how they experience withdrawal and which aspects of the human condition they most accurately model. These differences highlight each species’ strengths as model systems and can inform experimental design in studies of opioid withdrawal.

## Introduction

Opioid abuse is a devastating public health concern that has been amplified by the introduction of fentanyl and similar synthetic opioids as adulterants and primary drugs of abuse ^1^. The distinctly unpleasant signs and symptoms of opioid withdrawal, including increased sensitivity to pain (hyperalgesia), anxiety, gastrointestinal upset, lacrimation and rhinorrhea ^2^ hinder recovery and promote relapse. Compared to some but not all opioids, fentanyl has a higher affinity to opioid receptors and different effects on signaling bias ^3^ ^4^, reviewed by ^5–8^.

Hyperalgesia and lower pain tolerance during opioid withdrawal are prominently associated with increased relapse rates ^9–15^. As these symptoms can persist for extended periods after abstinence, and can appear as analgesic tolerance in the clinical setting ^16,17^, improving our understanding of the mechanisms underlying withdrawal, and pain in particular, will enable increasingly safe and effective support for patients who are attempting to discontinue opioid use.

Animal models are important for the pursuit of opioid withdrawal therapies, but elucidating the mechanisms driving specific withdrawal signs is complicated by conflicting results. This is particularly evident with respect to withdrawal induced hyperalgesia which has received relatively little attention. Some rodent models have demonstrated hyperalgesia to mechanical and thermal stimuli ^18–26^. Others find that hyperalgesia is absent or inconsistent ^27–30^. Furthermore, rodent studies rely almost exclusively on reflexive measures of thermal or mechanical sensitivity and do not assess measures of ongoing, spontaneous pain – a prominent feature of withdrawal in humans ^2^. Models that include both pain and somatic measures of withdrawal and that include both rats and mice to address species-specific differences have the potential to reduce conflicting reports by providing a more complete picture of opioid withdrawal and informing which species more accurately reflect specific aspects of the human withdrawal experience.

Here, we combine behavioral assays to assess both somatic withdrawal and pain related signs in fentanyl exposed rats and mice. We find that fentanyl withdrawal signs are largely similar to those frequently observed for morphine. However, mice and rats display unique patterns of somatic withdrawal behaviors and exhibit contrasting features of withdrawal-associated pain, suggesting that both species offer unique advantages in modeling withdrawal and associated pain.

## Methods

### Rigor

We adhered to accepted standards for rigorous study design and reporting to maximize the reproducibility and translational potential of our findings as described by Landis et al. ^31^ and in ARRIVE (Animal Research: Reporting In Vivo Experiments).

### Animals

All procedures were performed in accordance with the Animal Welfare Act regulations and Public Health Service guidelines and approved by the University of Maryland School of Medicine Animal Care and Use Committee. While opioid withdrawal is an inherently aversive condition, we selected the minimal exposure duration and lowest intensity stimuli for hyperalgesia testing. Noxious stimuli were immediately stopped after the first behavioral response. We monitored all animals throughout the study for signs of distress outside the immediate withdrawal testing period, including: continued weight loss, dehydration, lack of grooming, social isolation, excessive vocalization and fighting or injury. Of these signs, only minor weight loss occurred and was at a level consistent with prior studies of withdrawal ^32–34^. This weight loss is reported in detail in the results. We studied 29 Wistar Rats (saline: 6 males, 6 females; fentanyl: 7 males, 10 females) obtained from Charles River (Wilmington, MA), ranging in weight from 247 to 586 grams and 3 to 6 months of age at baseline. We studied 61 C57BL6/J mice (saline: 16 saline males, 16 females; fentanyl: 13 males, 16 females) obtained from Jackson Laboratory (Farmington, CT) and ranging in weight from 18 to 35 grams and 2 to 5 months of age at baseline.

### Drugs

Fentanyl Citrate (Cayman Chemicals, Ann Arbor, MI) and Naloxone Hydrochloride (Sigma, St. Louis, MO) were diluted in sterile saline and injected subcutaneously for all experiments at the concentrations indicated below.

### Acclimation to handling

Rats were acclimated to handling and to the experimental room for a minimum of 10 minutes per day, 5 days per week, for 2 weeks prior to beginning baseline assessments. Mice were acclimated to the experimental room for 30 minutes before testing.

### Fentanyl Exposure

We exposed animals to fentanyl or saline using subcutaneous (s.c.) injections twice daily (morning and afternoon) for a total of 9 exposure days across 2 weeks (excluding weekends, to mimic intermittent patterns of human use). We escalated the fentanyl dose over the first three days of exposure, divided into two separate injections (morning and afternoon dose), starting at 0.7 mg/kg twice daily and escalating to 1.3 mg/kg twice daily for mice and 0.04 to 0.10 mg/kg twice daily for rats. The total daily dose ranged from 1.4 to 2.6 mg/kg per day for mice and 0.08 to 0.20 mg/kg for rats. The animals were maintained at these doses for the remainder of the experiment. We selected these doses of fentanyl by designing the escalations to remain below the LD_50_ values for male mice of 62 mg/kg ^35^ and female rats of 2.91 mg/kg ^36^, while starting at a high enough initial dose to elicit opioid exposure signs in pilot animals. Suspected overdoses occurred in 2 rats and 3 mice, all males.

### Locomotor testing

Because hyperactivity is a hallmark of chronic exposure to opioids like morphine, a subset of mice (n=6/sex/drug) underwent locomotor testing during fentanyl exposure. After habituation to an open field, mice were injected with saline (10 mL/kg) to determine baseline locomotion over 30 min with video tracking software (CleverSys, Reston, VA). The following day, mice received a single s.c. injection of 2.5 mg/kg fentanyl or saline and were placed back in the open field for an additional 30 min. This process was repeated each day for a total of 5 fentanyl (or saline) injections.

### Withdrawal Paradigms

Figure 1 outlines the time-course of experiments. Spontaneous withdrawal was assessed 16 to 18 hours after the last injection, that is, the morning after the previous day’s afternoon dose. This was immediately followed by mechanical and thermal sensitivity testing performed at 20-22 hours of withdrawal. This spontaneous withdrawal testing timeframe was chosen based on pilot data that suggested qualitatively that somatic signs peak during this period (Fig. 2). After spontaneous withdrawal, animals were given the afternoon dose of fentanyl and returned to their home cages over the weekend. To evoke precipitated withdrawal, in the same animals that had previously undergone spontaneous withdrawal we administered one half of a total day’s dose of fentanyl (1.3 mg/kg for mice and 0.1mg/kg for rats), or an equal volume of saline for the control group. After 5 minutes we administered s.c. naloxone (10 mg/kg for mice, 3 mg/kg for rats, both fentanyl and saline groups), and performed withdrawal observations and testing during the following 30 to 60 minutes. As stated above, we chose fentanyl doses based on LD_50_ values for mice as compared to rats—values that reflect differences in metabolic rate for each species. Thus, we chose dosages of naloxone based on these differential doses of fentanyl, as well as pilot tests of naloxone-precipitated withdrawal with lower doses of naloxone (1mg/kg for rats, 1 mg/kg for mice) which yielded qualitatively mild behavioral responses.

**Figure 1:**
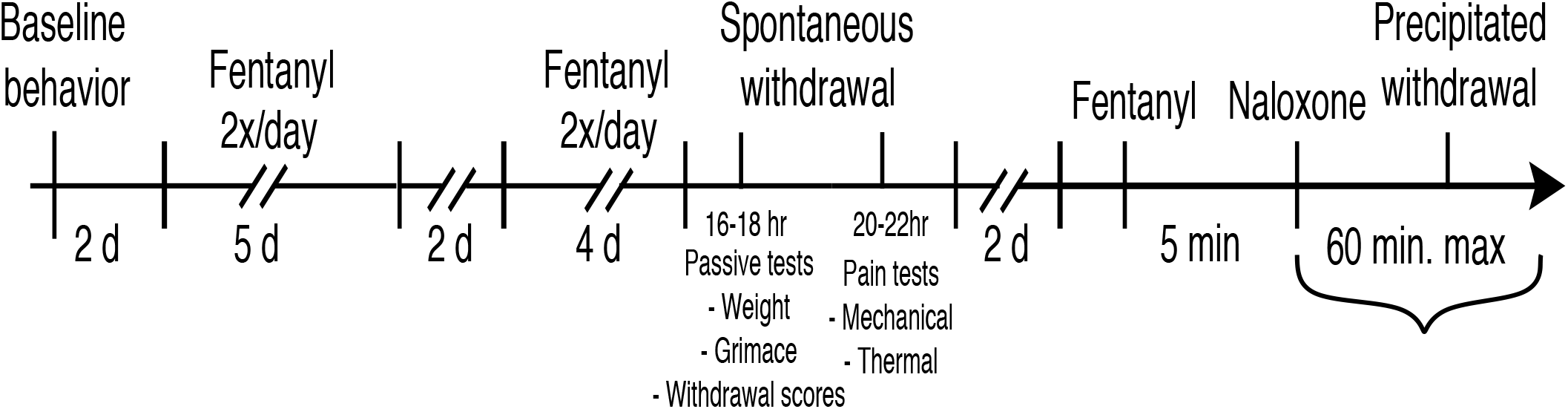
Timeline of drug administration and behavioral testing. We assessed baseline behavior on Fridays and gave animals the weekend off. We then began 2x daily fentanyl or saline administration, for 9 cumulative days, with weekends off. Spontaneous withdrawal was assessed the day after the last drug exposure. After 2 days without manipulation, we injected animals with fentanyl or saline, waited 5 minutes, then injected naloxone to assess precipitated withdrawal.

**Figure 2:**
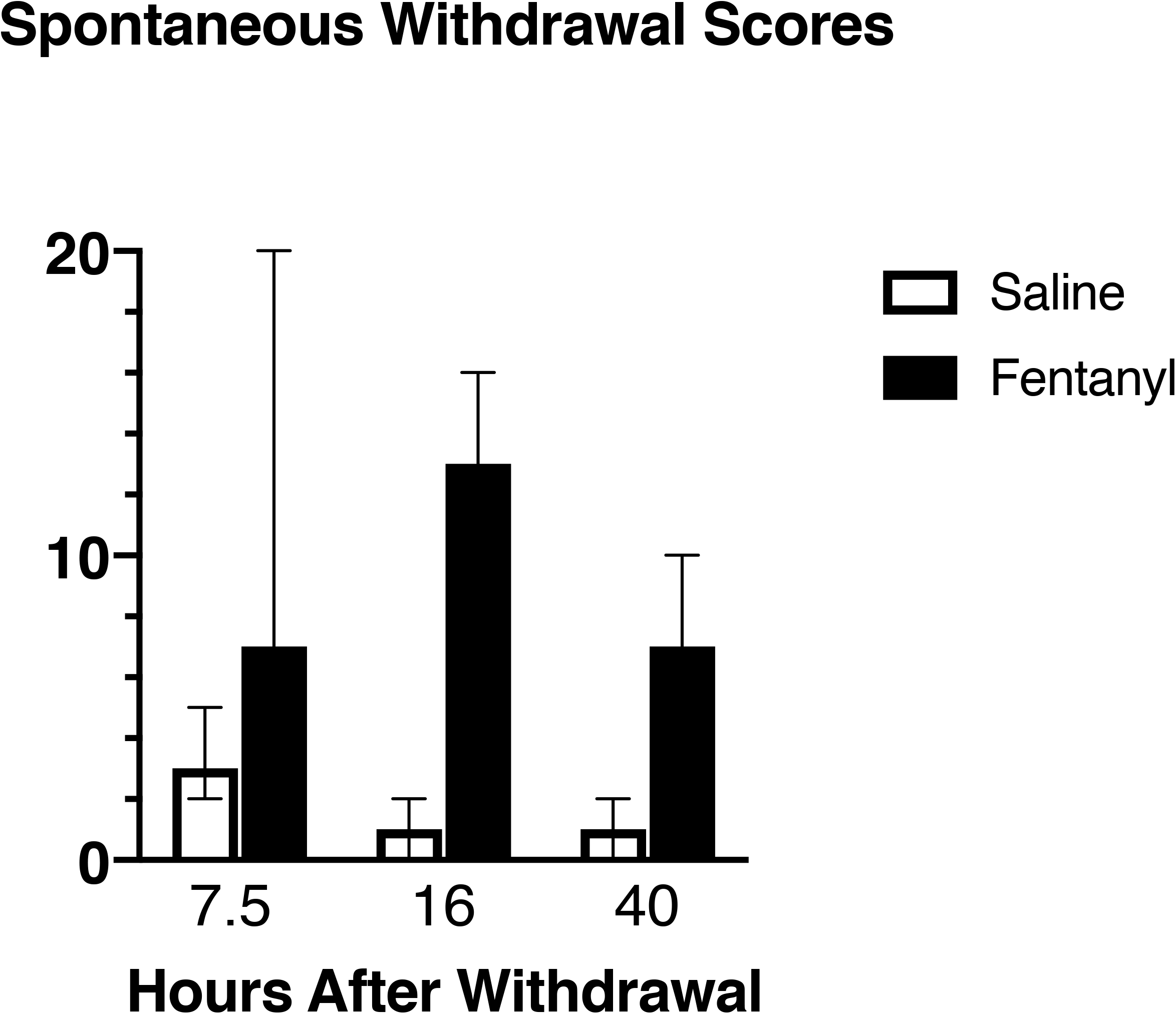
Timecourse of withdrawal scores during spontaneous withdrawal from subcutaneous fentanyl. Animals completed the 2-week intermittent, subcutaneous fentanyl (or saline) exposure protocol. For each animal, behavior was scored based on a 10-minute observation at 7.5, 16 and 40 hours after the final dose of fentanyl. Pilot data from n=4 saline treated rats and n=4 fentanyl treated rats. Data bars represent medians, with error bars showing 95% Confidence intervals.

### Withdrawal Scoring

The withdrawal behaviors we assessed were diarrhea, facial fasciculations/teeth chattering, abdominal constrictions/writhing, ptosis, wet dog shakes, abnormal posture, erection/ejaculation, genital grooming, excessive swallowing, hyperirritability, chromodacryorrhea (rats only), piloerection, persistent trembling (mice only) and paw tremor. We assessed body weight as an independent metric. We scored these behaviors using a modified version of the system developed by Gellert and Holtzman ^34^. Graded signs—a numerical score is assigned based on how many times a behavior occurred—included abdominal constrictions/writhing (2 points per writhe) and wet dog shakes (2 points for 1-2 shakes and 4 points for 3 or more shakes). The remaining observed behaviors were checked signs, given a weighted numerical score based on their presence or absence. For these checked signs, behaviors receiving a score of 2 points were: diarrhea, facial fasciculations/teeth chattering, ptosis, genital grooming, excessive swallowing, piloerection and paw tremor. Behaviors receiving a score of 3 points were: abnormal posture, erection/ejaculation, hyperirritability (defined as thrashing and vocalization upon handling, resisting placement both into and out of plexiglass chambers and home cage), and persistent trembling (defined as lasting more than 2 seconds). Chromodacryorrhea (rats only) received a score of 5 points. For spontaneous withdrawal, we observed rats in their home cages for 10 minutes. For naloxone precipitated withdrawal, we observed rats for precipitated withdrawal signs for 10 minutes in plexiglass containers, while obtaining grimace footage (see below). We chose this limited observation time window to maximize inclusion of additional tests before the effects of naloxone wore off. Naloxone has an average plasma half-life of approximately 30 minutes in rats ^37^ and 30 to 60 minutes in mice ^38^. Therefore, we observed rats for precipitated withdrawal signs for 10 minutes while they were in plexiglass chambers (8”x8”) undergoing filming for grimace analysis (see below). For mice, all withdrawal scoring was performed using 10-minute videos of mice in a plexiglass chamber (4” × 2” X 2.5”).

### Body Weights

For rats, we gathered weights prior to baseline behavior and every morning prior to initial fentanyl injections, as well as immediately before spontaneous withdrawal behavioral testing. On the precipitated withdrawal testing day, we weighed rats immediately before the fentanyl injection and at the close of all testing (maximum of 60 minutes after the naloxone injection). For mice, we gathered weights before baseline behavior, every morning prior to initial fentanyl injection and after spontaneous withdrawal behavioral testing. We did not collect mouse weights on precipitated withdrawal testing days.

### Fecal Boli

For naloxone-precipitated withdrawal in rats only, we counted the number of fecal boli produced in the precipitated withdrawal time frame and reported this as a separate metric. We did not quantify boli in rats in spontaneous withdrawal, or in mice in either spontaneous or precipitated withdrawal because the apparatus contained a wire mesh floor making it incompatible with this metric.

### Mechanical Sensitivity

For rats, to assess mechanical sensitivity we applied calibrated von Frey filaments (North Coast Medical, Gilray, CA) ranging from 8 to 300 grams to the plantar surface of the right hindpaw. We defined a response as a sharp paw withdrawal, licking, or shaking, and used the up-down method across 10 trials ^39^ to determine each animal’s withdrawal threshold. Trials where the response pattern did not converge to a stable value were discarded. For mice, we applied a single supra-threshold filament of 2.0 grams to the plantar surface of the hindpaw 3 times and scored responses on a scale of 1 (no response or foot passively pushed up by pressure of filament) to 4 (repeated shaking or prolonged lift or licking for more than 5 seconds). From these results we generated a binary scoring system. Animals that received a score of 0 to 2 received a binary score of 1. Animals that received a score of 3 to 4 received a binary score of 2. The scores from the 3 trials were then averaged.

### Thermal Sensitivity (Heat)

For rats, we used the Hargreaves apparatus (IITC, Woodland Hills, CA) over 3 trials to assess heat sensitivity, by aiming the light beam at the plantar surface of the right hindpaw and recording the latency to paw withdrawal. We use the median value of the 3 trials as the reported value for each rat, and then calculated percent of baseline latencies during spontaneous and precipitated withdrawal. For mice, we used the hotplate test. Mice were covered by a small beaker on top of the hotplate (Hot Plate Analgesia Meter for Mice and Rats Series 8, IITC Life Science) and the temperature was gradually increased from 33 to 50°C over a 2-minute period. The time it took for mice to respond (grooming/licking hind paw, attempt to escape, etc.) was reported.

### Thermal Sensitivity (Cold)

To assess cold sensitivity, we used a 1 mL syringe to apply 1 drop of acetone (Fisher Scientific, Hampton, NH) to the plantar surface of the hindpaw. We performed 3 trials of acetone application at least 5 minutes apart. Scores were then obtained, the minimum score being 0 (no response to any of the 3 trials) and the maximum possible score being 3 (repeated flicking and licking of paws on each of the 3 trials). From these results, we then generated a binary scoring system. Animals that received a score of 0 to 1 received a binary score of 1. Animals that received a score of 2 to 3 received a binary score of 2. In rats, acetone did not elicit a response in the saline or fentanyl groups from the initial cohorts and the test was discontinued for subsequent cohorts.

### Grimace Scoring

To assess ongoing, spontaneous pain in mice and rats, we used the grimace scoring system ^40–42^. On baseline and spontaneous withdrawal days, we filmed rats in plexiglass chambers for 30 minutes. Due to the time constraint of naloxone-precipitated withdrawal, we shortened the filming duration to 10 minutes to allow sufficient time to perform other behavioral testing. For mice, we recorded somatic spontaneous and precipitated behaviors for 10 minutes in plexiglass chambers prior to any other behavioral tests. These videos were also used for grimace analysis. We obtained 2 images per minute by capturing video screenshots. These images were then uploaded into MATLAB. For rats, we used the FaceFinder application ^41^, generously shared with us by J. Mogil, to select still images from video footage for action unit analysis. Presumably because of the darker fur and eyes of the C57Bl6/J mice, we manually selected still images for this species since FaceFinder did not successfully capture usable images. This discrepancy did not affect the scoring itself, since at the end of both processes, we were left with ten clear images selected from each animal, at each timepoint, for analysis. According to prior reports ^41,42^ we scored each action unit on a scale from 0 (minimal pain) to 2 (multiple signs of pain) for each image. We used a custom-written MATLAB script (Natick, MA) which allowed all images from all timepoints to be scrambled so that the scorer was blind to timepoint and experimental group. This MATLAB script then averaged each image’s score to calculate a “grand average” grimace score for each animal at each timepoint.

### Statistical Analysis

We present grimace scores and withdrawal scores in their raw form. For heat sensitivity, cold sensitivity, mechanical sensitivity, and body weights, post-exposure fentanyl withdrawal data are presented as a percent of baseline, to facilitate comparisons between animals and between species. For reference, raw values for each metric are provided in Table 1 for baseline, Table 2 for spontaneous and Table 3 for naloxone precipitated withdrawal. We analyzed all data and generated figures using GraphPad PRISM version 8.2.0 for Mac (GraphPad Software, La Jolla, CA). We used nonparametric statistical tests where the necessary criteria to use parametric tests were not met (normal distribution as assessed by D’agostino-Pearson test, independent data, and homogenous variance). If data were normally distributed but had unequal variances (as assessed by PRISM’s F-test to compare variances), we use a Welch’s t-test.

Although sample sizes were not sufficient to fully assess sex differences, we ran a 2-way ANOVA with sex and treatment (fentanyl or saline) as variables: no sex differences were observed in any metric.

**Table 1:** Baseline Behaviors. Raw data values at baseline. Values shown are median with 95% CI, followed by minimum, maximum, standard deviation, and sample size. Note that we assessed baseline behavior prior to any drug exposure, so sample sizes represent animals from eventual saline and fentanyl groups.

| Test | Value | Mice | Rats |
| --- | --- | --- | --- |
| Withdrawal Scores | Median ± 95% CI | 4, 1-5 | 2, 0-5 |
|  | Min | 0 | 0 |
|  | Max | 22 | 9 |
|  | SD | 4.7 | 2.7 |
|  | n | 27 | 14 |
| Mechanical (score for mice, grams for rats) | Median ± 95% CI | 1.0, 1.0- 1.0 | 37.5, 29.9-45.1 |
|  | Min | 1 | 12.3 |
|  | Max | 1.3 | 134.9 |
|  | SD | 0.09 | 28.9 |
|  | n | 37 | 25 |
| Heat<br>(hotplate or Hargreaves, seconds) | Median ± 95% CI | 102.5, 96.3-107.1 | 4.1, 3.4-5.0 |
|  | Min | 72.4 | 2.2 |
|  | Max | 122 | 6.5 |
|  | SD | 12.2 | 1.1 |
|  | n | 37 | 21 |
| Cold | Median ± 95% CI | 1.0, 1.0-1.3 | n/a, did not respond |
|  | Min | 1.0 | n/a |
|  | Max | 1.67 | n/a |
|  | SD | 0.25 | n/a |
|  | n | 37 | 16 |
| Grimace | Median ± 95% CI | 0.28, 0.17-0.35 | 0.5, 0.35-0.68 |
|  | Min | 0 | 0.09 |
|  | Max | 0.88 | 1.1 |
|  | SD | 0.17 | 0.27 |
|  | n | 27 | 29 |

**Table 2:**
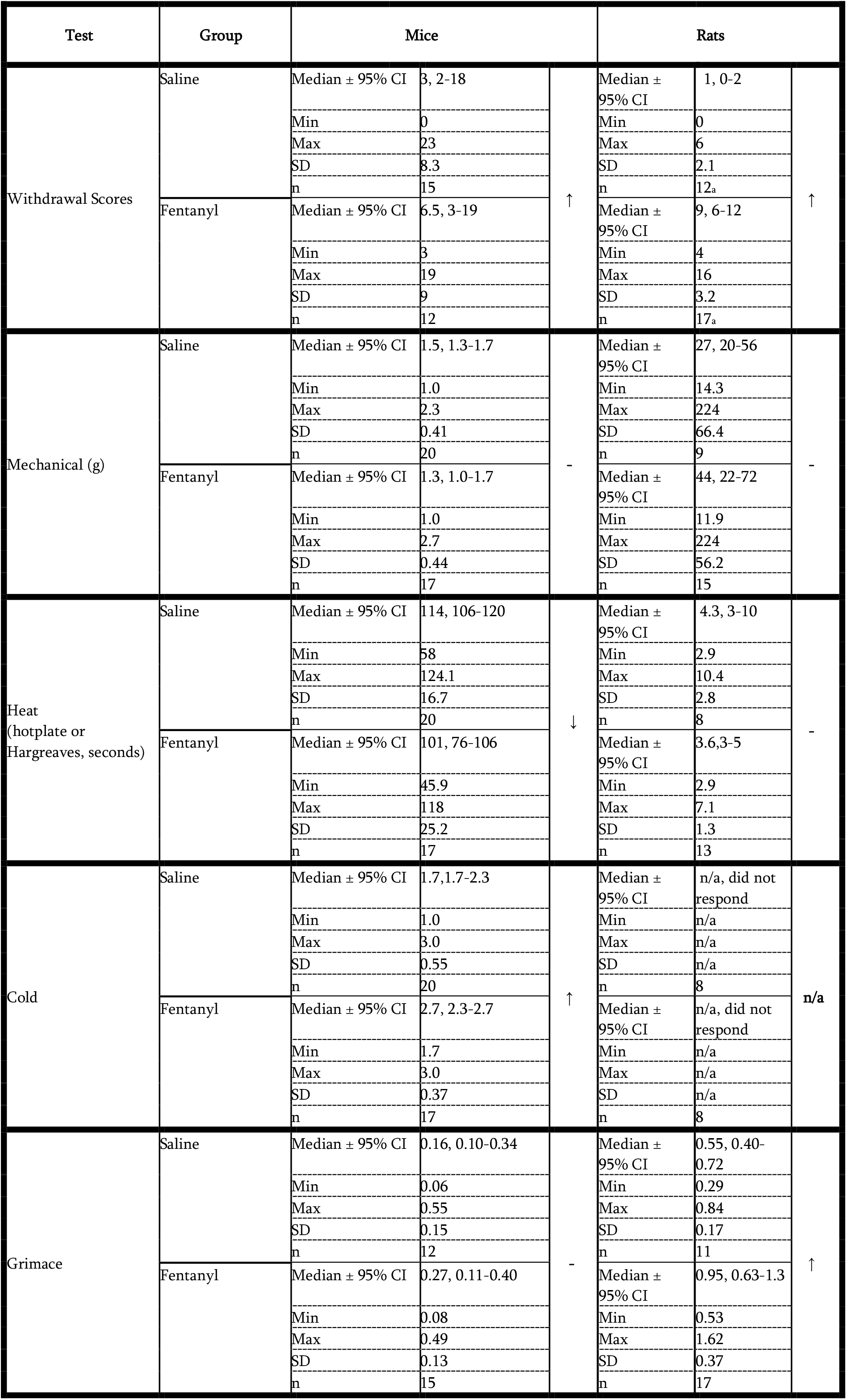
Spontaneous Withdrawal Behaviors. Raw data values and summary of behavioral changes in spontaneous withdrawal. Note that higher acetone and grimace scores, but lower heat withdrawal latencies indicate more pain. Arrows indicate change in fentanyl relative to saline groups. Values shown are median with 95% CI, followed by minimum, maximum, standard deviation, and sample size.

**Table 3:**
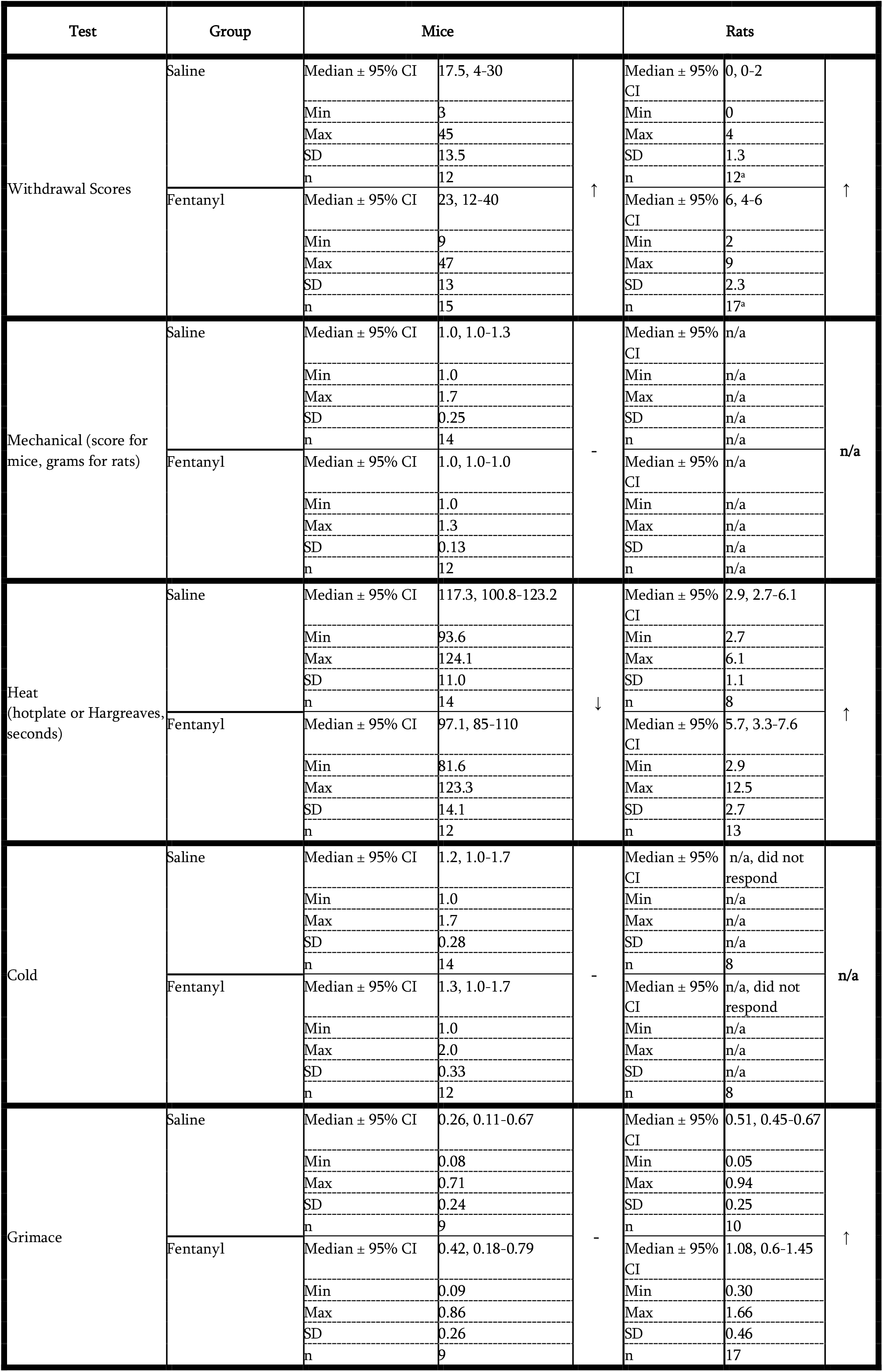
Raw data values and summary of behavioral changes in naloxone precipitated withdrawal. Note that higher acetone and grimace scores, but lower heat withdrawal latencies indicate more pain. Arrows indicate change in fentanyl relative to saline groups. Values shown are median with 95% CI, followed by minimum, maximum, standard deviation, and sample size.

| Test | Group | Mice |  |  | Rats |  |  |
| --- | --- | --- | --- | --- | --- | --- | --- |
| Withdrawal Scores | Saline | Median ± 95% CI | 17.5, 4-30 | ↑ | Median ± 95% CI | 0, 0-2 | ↑ |
|  |  | Min | 3 |  | Min | 0 |  |
|  |  | Max | 45 |  | Max | 4 |  |
|  |  | SD | 13.5 |  | SD | 1.3 |  |
|  |  | n | 12 |  | n | 12<sup>a</sup> |  |
|  | Fentanyl | Median ± 95% CI | 23, 12-40 | ↑ | Median ± 95% CI | 6, 4-6 | ↑ |
|  |  | Min | 9 |  | Min | 2 |  |
|  |  | Max | 47 |  | Max | 9 |  |
|  |  | SD | 13 |  | SD | 2.3 |  |
|  |  | n | 15 |  | n | 17<sup>a</sup> |  |
| Mechanical (score for mice, grams for rats) | Saline | Median ± 95% CI | 1.0, 1.0-1.3 | - | Median ± 95% CI | n/a | n/a |
|  |  | Min | 1.0 |  | Min | n/a |  |
|  |  | Max | 1.7 |  | Max | n/a |  |
|  |  | SD | 0.25 |  | SD | n/a |  |
|  |  | n | 14 |  | n | n/a |  |
|  | Fentanyl | Median ± 95% CI | 1.0, 1.0-1.0 | - | Median ± 95% CI | n/a | n/a |
|  |  | Min | 1.0 |  | Min | n/a |  |
|  |  | Max | 1.3 |  | Max | n/a |  |
|  |  | SD | 0.13 |  | SD | n/a |  |
|  |  | n | 12 |  | n | n/a |  |
| Heat (hotplate or Hargreaves, seconds) | Saline | Median ± 95% CI | 117.3, 100.8-123.2 | ↓ | Median ± 95% CI | 2.9, 2.7-6.1 | ↑ |
|  |  | Min | 93.6 |  | Min | 2.7 |  |
|  |  | Max | 124.1 |  | Max | 6.1 |  |
|  |  | SD | 11.0 |  | SD | 1.1 |  |
|  |  | n | 14 |  | n | 8 |  |
|  | Fentanyl | Median ± 95% CI | 97.1, 85-110 | ↓ | Median ± 95% CI | 5.7, 3.3-7.6 | ↑ |
|  |  | Min | 81.6 |  | Min | 2.9 |  |
|  |  | Max | 123.3 |  | Max | 12.5 |  |
|  |  | SD | 14.1 |  | SD | 2.7 |  |
|  |  | n | 12 |  | n | 13 |  |
| Cold | Saline | Median ± 95% CI | 1.2, 1.0-1.7 | - | Median ± 95% CI | n/a, did not respond | n/a |
|  |  | Min | 1.0 |  | Min | n/a |  |
|  |  | Max | 1.7 |  | Max | n/a |  |
|  |  | SD | 0.28 |  | SD | n/a |  |
|  |  | n | 14 |  | n | 8 |  |
|  | Fentanyl | Median ± 95% CI | 1.3, 1.0-1.7 | - | Median ± 95% CI | n/a, did not respond | n/a |
|  |  | Min | 1.0 |  | Min | n/a |  |
|  |  | Max | 2.0 |  | Max | n/a |  |
|  |  | SD | 0.33 |  | SD | n/a |  |
|  |  | n | 12 |  | n | 8 |  |
| Grimace | Saline | Median ± 95% CI | 0.26, 0.11-0.67 | - | Median ± 95% CI | 0.51, 0.45-0.67 | ↑ |
|  |  | Min | 0.08 |  | Min | 0.05 |  |
|  |  | Max | 0.71 |  | Max | 0.94 |  |
|  |  | SD | 0.24 |  | SD | 0.25 |  |
|  |  | n | 9 |  | n | 10 |  |
|  | Fentanyl | Median ± 95% CI | 0.42, 0.18-0.79 | - | Median ± 95% CI | 1.08, 0.6-1.45 | ↑ |
|  |  | Min | 0.09 |  | Min | 0.30 |  |
|  |  | Max | 0.86 |  | Max | 1.66 |  |
|  |  | SD | 0.26 |  | SD | 0.46 |  |
|  |  | n | 9 |  | n | 17 |  |

## Results

### Mice and rats exhibit somatic withdrawal signs at baseline

Because animals can exhibit some withdrawal-associated behaviors in the absence of drug (e.g. genital grooming), we first characterized these behaviors at baseline (Table 1; Raw withdrawal scores for spontaneous and precipitated withdrawal are in Tables 2 and 3). Even before any drug exposure, mice displayed behaviors associated with withdrawal, with considerable variation between animals (n=27, median withdrawal score=4, 95% C.I. 1-5). Behaviors present in mice at baseline included grooming, genital grooming, teeth chattering, swallowing, persistent trembling, paw tremors, piloerection, and wet dog shakes. Rats also displayed somatic withdrawal behaviors at baseline (n=14, median withdrawal score = 2, 95% C.I. 0-5). Behaviors present in rats at baseline included teeth chattering, swallowing, ptosis, hyperirritability, and wet dog shakes.

### Mice and rats exhibit species-specific responses to acute fentanyl

After baseline behavioral assessments, we proceeded with drug exposure and qualitatively assessed animals’ responses to fentanyl (Timeline in Fig.1). These observations revealed consistent reactions within each species, despite notable differences between mice and rats. We chose fentanyl doses based on the LD_50_ for each species to account for differences in metabolic rate between species – mice have a more than 10-fold higher LD_50_ for fentanyl citrate administered subcutaneously, compared to rats (Cayman Chemicals). Within minutes after fentanyl injection, mice became hyperactive, moving swiftly and persistently around the cage perimeter, despite apparent muscle stiffness, including raised tails (Straub tail) and chest wall rigidity (barrel chested in appearance), consistent with previous reports in opioid-exposed mice ^43^. These signs gradually declined over the course of one to two hours, with the mice appearing normal within 3 hours. We saw no signs of adaptation to fentanyl exposure throughout the two-week period, and mice gradually developed hyperlocomotion. We quantitated locomotor activity over 5 days of drug or saline exposure. Fentanyl exposed mice ran increasingly longer distances across days, compared to saline-treated mice in an open field test (Fig. 3; Saline n = 12, Fentanyl n = 12, 2-way repeated-measures ANOVA. Effect of treatment: F(1,22) = 32.2, p < 10^−4^ Effect of day: F(5,110) = 17.4, p< 10^−4^. Effect of interaction between day and treatment: F(5,110) = 10.3, p < 10^−4^).

**Figure 3:**
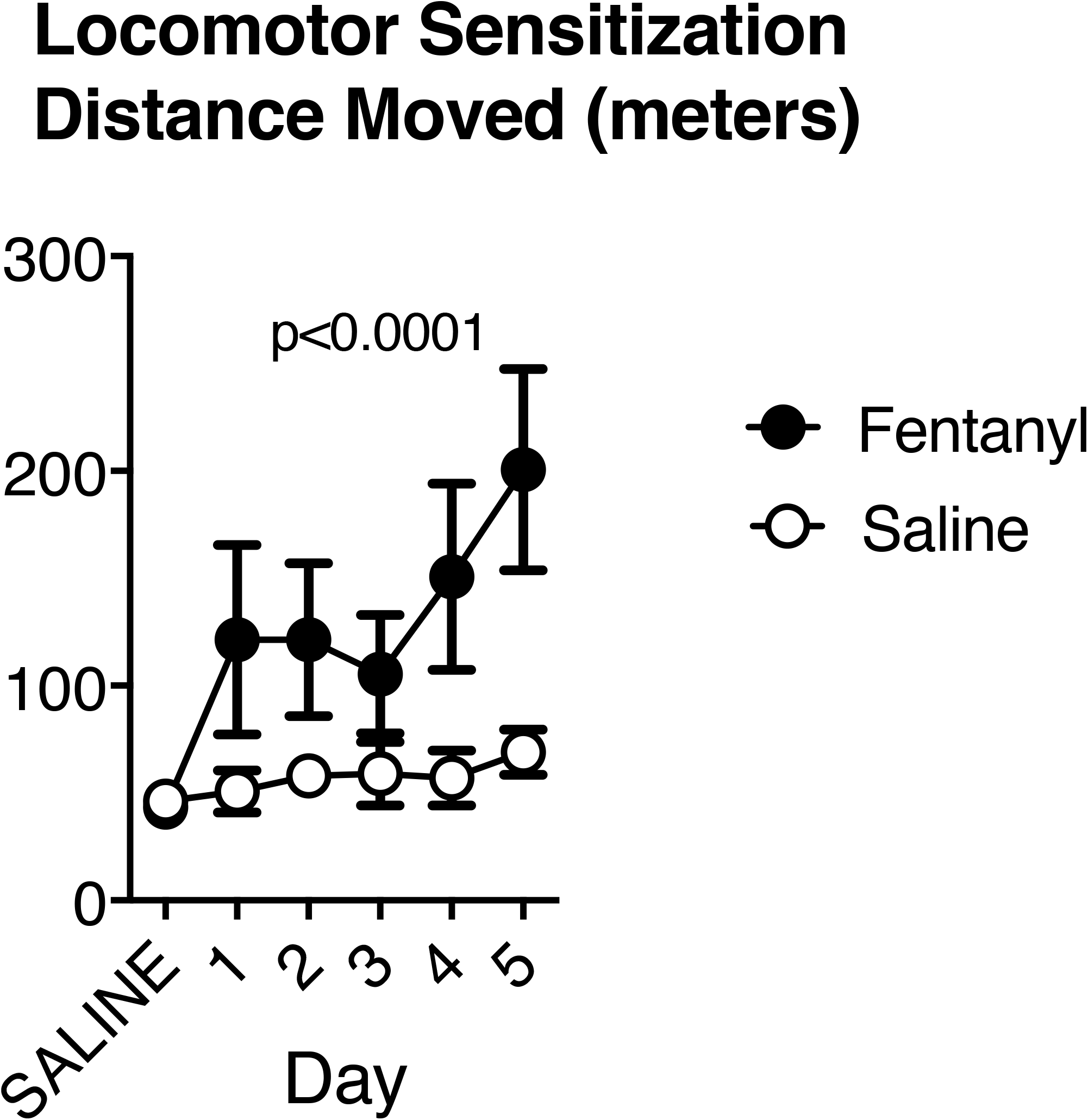
Mice develop hyperlocomotion in response to repeated fentanyl exposure. Across 5 days of subcutaneous injections of fentanyl (n=12 animals; 2.5mg/kg/day), mice traveled increasingly large total distances in an open field apparatus, compared to controls (n=12). Data represent means and 95% confidence intervals.

In contrast to mice, rats became lethargic and remained stationary, with exophthalmos and reduced respiratory rate. These symptoms persisted for approximately 30 minutes, before the rats gradually began moving around the cage. At this point, rats typically filled their mouths with bedding granules and started chewing. During week 2 of exposure, rats began to display oral stereotypy (gentle forelimb chewing) and hyperactivity, consistent with other reports of rat responses to chronic opioid exposure ^44–46^. This hyperactivity included short bursts of rapid locomotion from one side of the cage to the other (along the cage’s length), lasting up to 3 hours post-exposure.

### Spontaneous withdrawal signs differ in mice and rats

Our goal was to evaluate somatic and pain-related withdrawal behaviors in mice and rats during opioid withdrawal. Therefore, after establishing baseline behaviors and completing an intermittent fentanyl exposure protocol, we evaluated behavior after halting drug exposure (spontaneous withdrawal).

#### Weight Changes

Fentanyl exposed mice had greater reductions in weight on the last administration day, compared to saline exposed mice (Fig. 4A; Saline: n = 15, mean 100.6% of baseline, 95% C.I. 97.7 - 103.5%. Fentanyl: n = 12, mean 95.6%, 95% C.I. 91.9 - 99.2%. Cohen’s d = 0.93, t(25) = 2.41, p=0.029, unpaired t test). We calculated each rat’s weight during spontaneous withdrawal (16-24 hours since last fentanyl exposure) as a percent of their pre-exposure, baseline weight. Fentanyl exposed rats lost weight over the course of the experiments and, compared to saline exposed rats, had greater weight reductions at spontaneous withdrawal (Fig. 4A, Saline: n = 12: mean 103% of baseline, 95% C.I. 101 - 105%. Fentanyl: n = 17: mean 98% of baseline, 95% C.I. 96.9 - 99.2 %. Cohen’s d = 1.96, t(27) = 5.2, p < 10^−4^, unpaired t test.). There were no sex differences in percent of baseline weights or in any of the other metrics described, although larger sample sizes are needed to conclusively determine sex differences.

**Figure 4:**
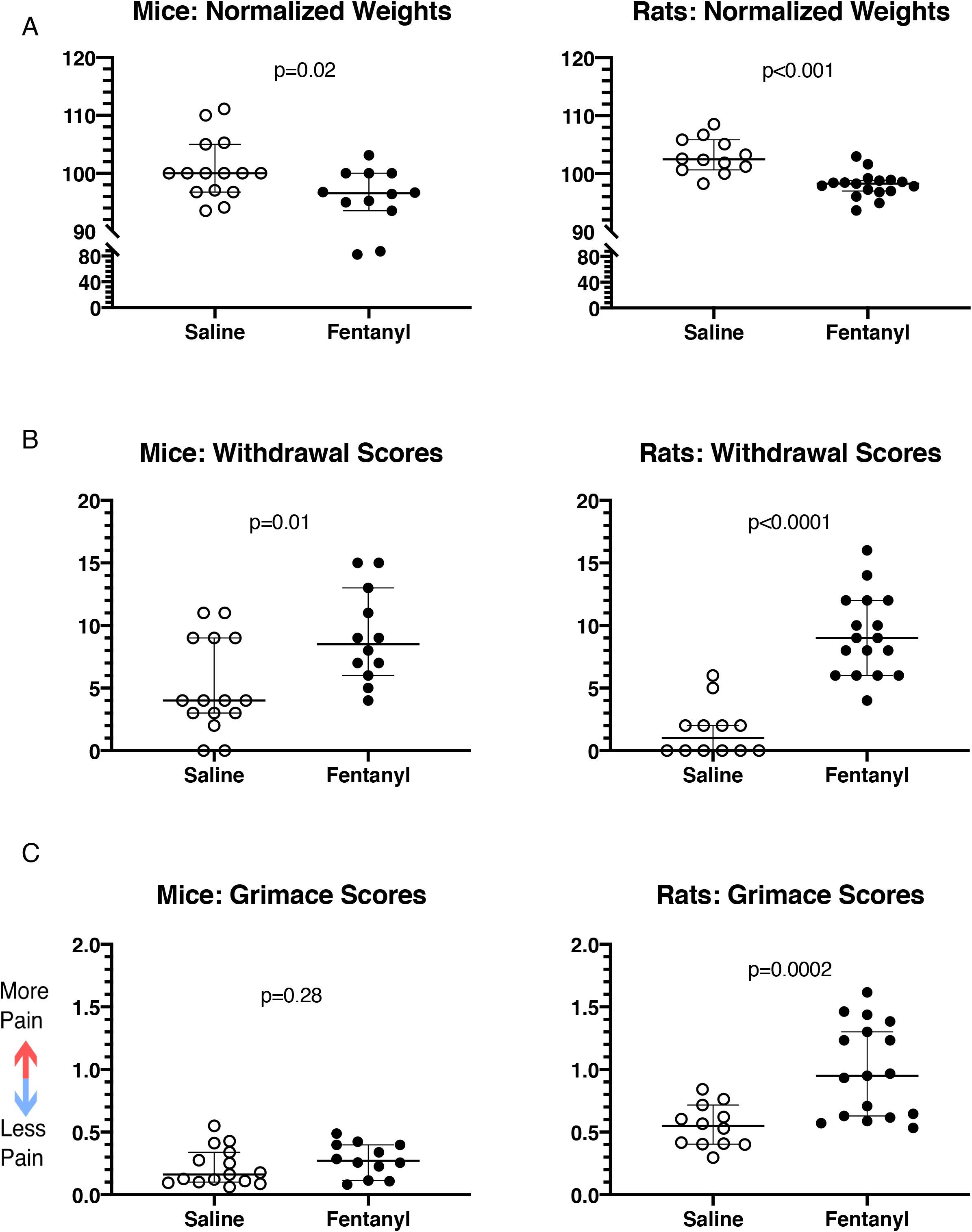
Passive behaviors in mice and rats during spontaneous fentanyl withdrawal. Fentanyl exposed mice and rats lose weight (A) and show more signs of withdrawal (B) relative to saline controls. However, only rats showed an increase in spontaneous pain metrics (C). Individual points represent data from a single animal, and horizontal lines depict medians and 95% confidence intervals. Normalized data are expressed as a percent of baseline.

#### Withdrawal Scores

Following the two-week exposure protocol (timeline in Fig. 1) we observed each animal for 10 minutes during the 16 to 18-hour spontaneous withdrawal time frame, and scored their withdrawal-associated behaviors, as described in Methods. Both mice and rats exposed to fentanyl showed higher withdrawal scores, compared to saline exposed animals (Fig. 4B). Fentanyl exposed mice (n = 12) had mean spontaneous withdrawal scores of 9.1 (95% C.I. 6.7 - 11.4), whereas saline exposed mice (n = 15) had a mean withdrawal score of 5.1 (95% C.I. 3.0 - 7.1, Cohen’s d = 1.09, t(25) = 2.8, p = 0.01, unpaired t test). Fentanyl exposed rats (n = 17) had mean withdrawal scores of 9.2 (95% C.I. 7.5 - 10.8, Cohen’s d = 1.09), whereas saline exposed rats (n = 12) had mean scores at spontaneous withdrawal of 1.6 (95% C.I. 0.27 - 2.9, Cohen’s d = 2.71, t(27) = 7.16, p < 10^−4^, unpaired t test).

Both saline and fentanyl treated mice exhibited multiple somatic withdrawal behaviors during the assessment period. However, a greater proportion of mice undergoing spontaneous fentanyl withdrawal exhibited wet dog shakes and piloerection (Fig 5A). In contrast, most somatic withdrawal behaviors were restricted to fentanyl treated rats, with few saline treated rats showing somatic withdrawal behaviors.

**Figure 5:**
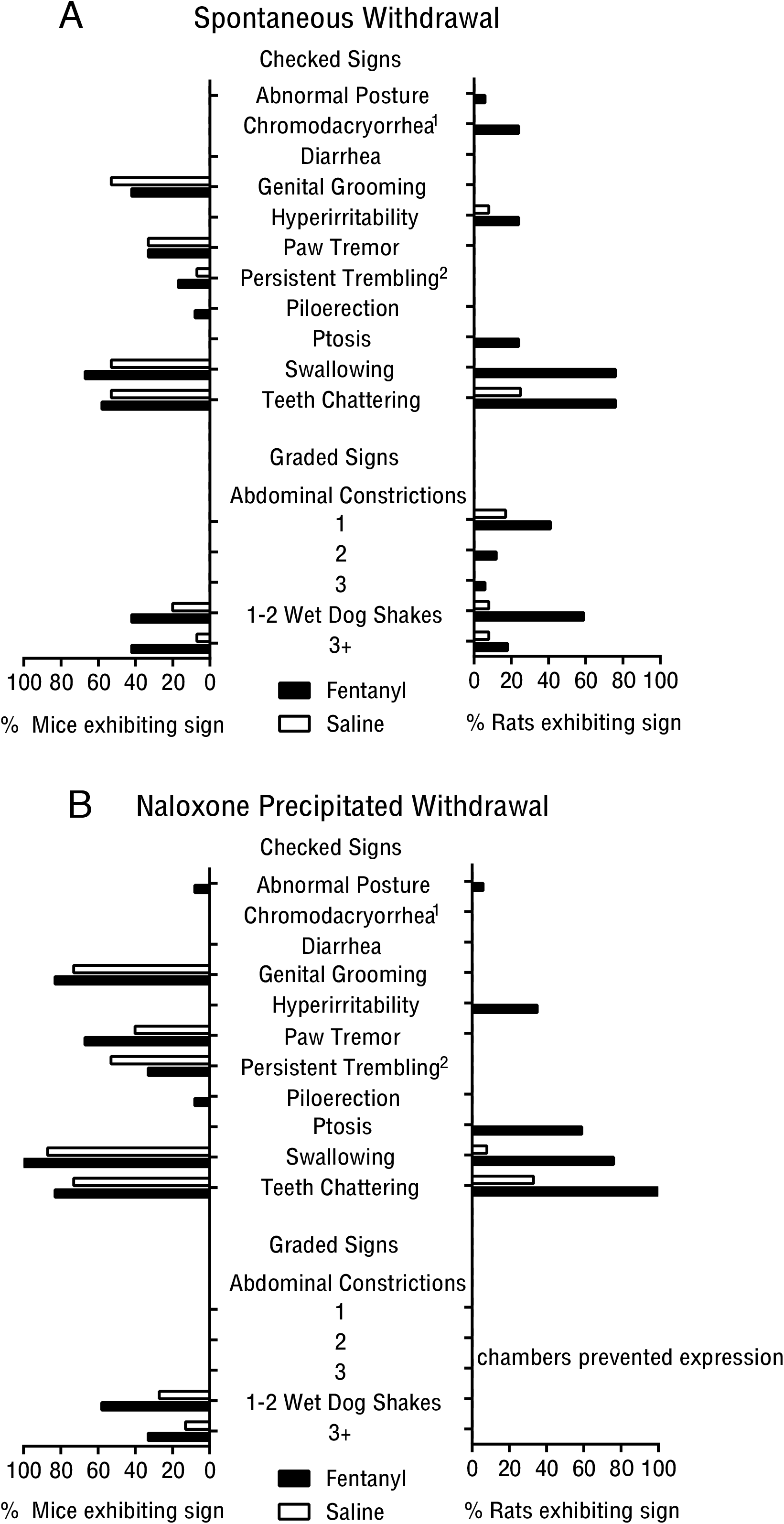
Prevalence of somatic withdrawal signs in mice and rats during (A) spontaneous and (B) naloxone precipitated fentanyl withdrawal. Superscripts denote behaviors assessed exclusively in rat(1) or mouse(2). All other data points of 0 reflect an absence of behavior, not absence of observation.

### Mice show thermal hypersensitivity and rats show increased ongoing pain during spontaneous withdrawal

#### Ongoing Pain

To evaluate ongoing pain, we assessed grimace scores, a well-validated metric for spontaneous pain in rodents ^40–42^. Grimace scores of fentanyl exposed mice during spontaneous withdrawal did not differ from those in saline treated mice (Fig 4C; fentanyl: n = 12, mean score. 0.28, 95% C.I. 0.20 - 0.37; saline: n = 15, mean score 0.22, 95% C.I. 0.14 - 0.30; t(25) = 1.1, p = 0.28, unpaired t test). Fentanyl exposed rats had higher grimace scores during spontaneous withdrawal, compared to saline exposed rats, suggesting that they were in more pain (Fig. 4C; saline: n = 12, mean score 0.54, 95% C.I. 0.44 - 0.65; fentanyl: n = 17, mean score 0.99, 95% C.I. 0.8 - 1.2. Cohen’s d = 1.44, t(23.7) = 4.3, p = 0.0002, Welch’s t-test).

#### Thermal Sensitivity (Heat)

Mice and rats displayed different patterns of heat sensitivity 20-22 hours into spontaneous withdrawal. Fentanyl exposed mice had shorter-latency hot plate responses (Fig. 6A), indicative of increased thermal sensitivity (n=17, median 94.1% of baseline, 95% C.I. 82.3-101.7%), compared to saline exposed mice (n=20, median 109.6% of baseline, 95% C.I. 101.9-117.1%; Cohen’s d=0.82, Mann-Whitney U=79, p=0.0048), suggesting that they developed heat hypersensitivity during this time frame of withdrawal. Fentanyl exposed rats (n = 13, mean 109.8%, 95% C.I. 91.3- 128.3%) and saline exposed rats (n = 8, mean 121.7%, 95% C.I. 68.6 - 174.8%) had indistinguishable hind paw heat withdrawal latencies at 20 - 22 hours of spontaneous fentanyl withdrawal (Fig. 6A; t(9.04) = 0.5, p = 0.63, Welch’s t-test), suggesting that heat sensitivity was unaffected in rats.

**Figure 6:**
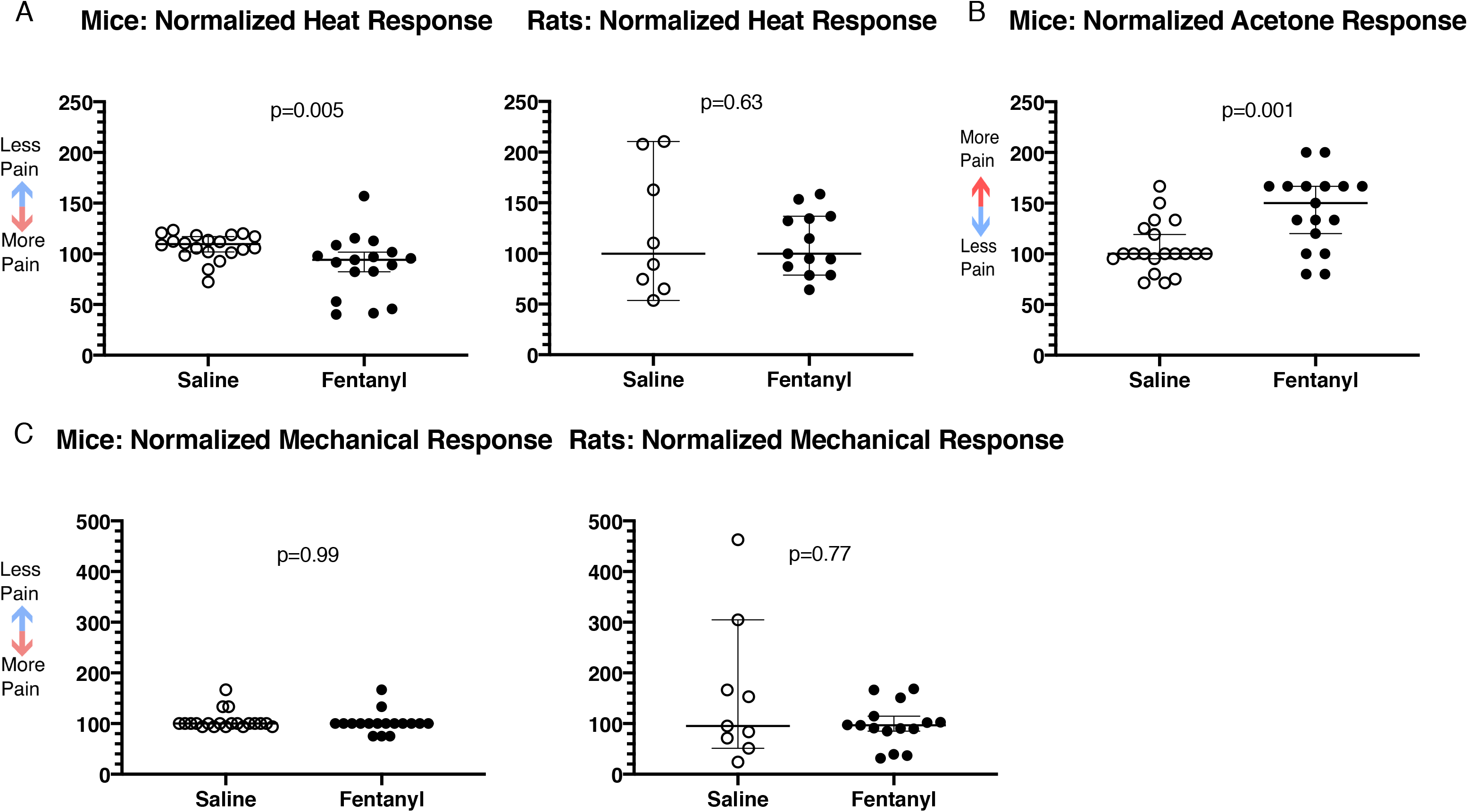
Pain related behaviors during spontaneous fentanyl withdrawal. (A) Mice show increased heat sensitivity using the hotplate test, whereas rats exhibit no change using the Hargreaves test. (B) Mice have increased cold sensitivity during spontaneous fentanyl withdrawal. (C) Mice and rats show no change in mechanical sensitivity. Individual points represent data from a single animal, and horizontal lines depict medians and 95% confidence intervals. Normalized data are expressed as a percent of baseline.

#### Thermal Sensitivity (Cold)

Fentanyl exposed mice had increased response scores to acetone at 20 - 22 hours spontaneous withdrawal (Fig. 6B; n = 17, mean 142.9%, 95% C.I. 123.7 - 162.2%), compared to saline exposed mice (n = 20, mean 105.8%, 95% C.I. 93.9 - 117.6%; Cohen’s d = 1.2, t(35) = 3.59, p = 10^−3^, unpaired t test). In rats, acetone failed to evoke any responses, in either experimental group (saline: n = 8; fentanyl: n = 8, data not shown).

#### Mechanical Sensitivity

Since touch hypersensitivity (mechanical allodynia) can occur in opioid withdrawal, we tested for changes in tactile sensitivity by quantifying animals’ responses to mechanical stimuli of the hindpaw. Mechanical sensitivity was unaffected at 20 - 22 hours spontaneous withdrawal in mice and rats, suggesting that they did not develop mechanical hypersensitivity (Fig. 6C; Mice: Mann-Whitney U=169.5, p=0.99. Rats: Mann-Whitney U=62, p=0.77). Thresholds in saline exposed (n=20) and fentanyl exposed (n=17) mice were 100% of their baseline values (95% C.I. confined to 100%). Similarly, thresholds in saline exposed rats (n=9) were 95% of their baseline values (95% C.I. 51.4-304.6%), and in fentanyl exposed rats (n=15) they were 96.5% of baseline (95% C.I. 84.9-114.6%).

### Naloxone-precipitated withdrawal signs differ between mice and rats

Because naloxone-precipitated withdrawal offers precise temporal control over symptom onset, we next determined profiles of somatic and pain-associated behaviors by administering naloxone 5 min after a fentanyl challenge (timeline in Fig 1).

#### Weight Changes

Fentanyl exposed rats had greater weight reductions after naloxone administration, compared to saline exposed rats (Fig. 7A; Saline: n=12, median 100.2% of pre-naloxone weight, 95% C.I. 100-100.3%. Fentanyl: n=17, median 99.62%, 95% C.I. 99-99.7%. Cohen’s d=1.48, Mann-Whitney U=21, p=10^−4^). In rats, we recorded weight immediately before fentanyl and naloxone injection, and again after completion of naloxone-precipitated withdrawal testing (see Methods). As with spontaneous withdrawal, there were no sex differences in weight changes. We did not record mouse weights before or after naloxone-precipitated withdrawal.

**Figure 7:**
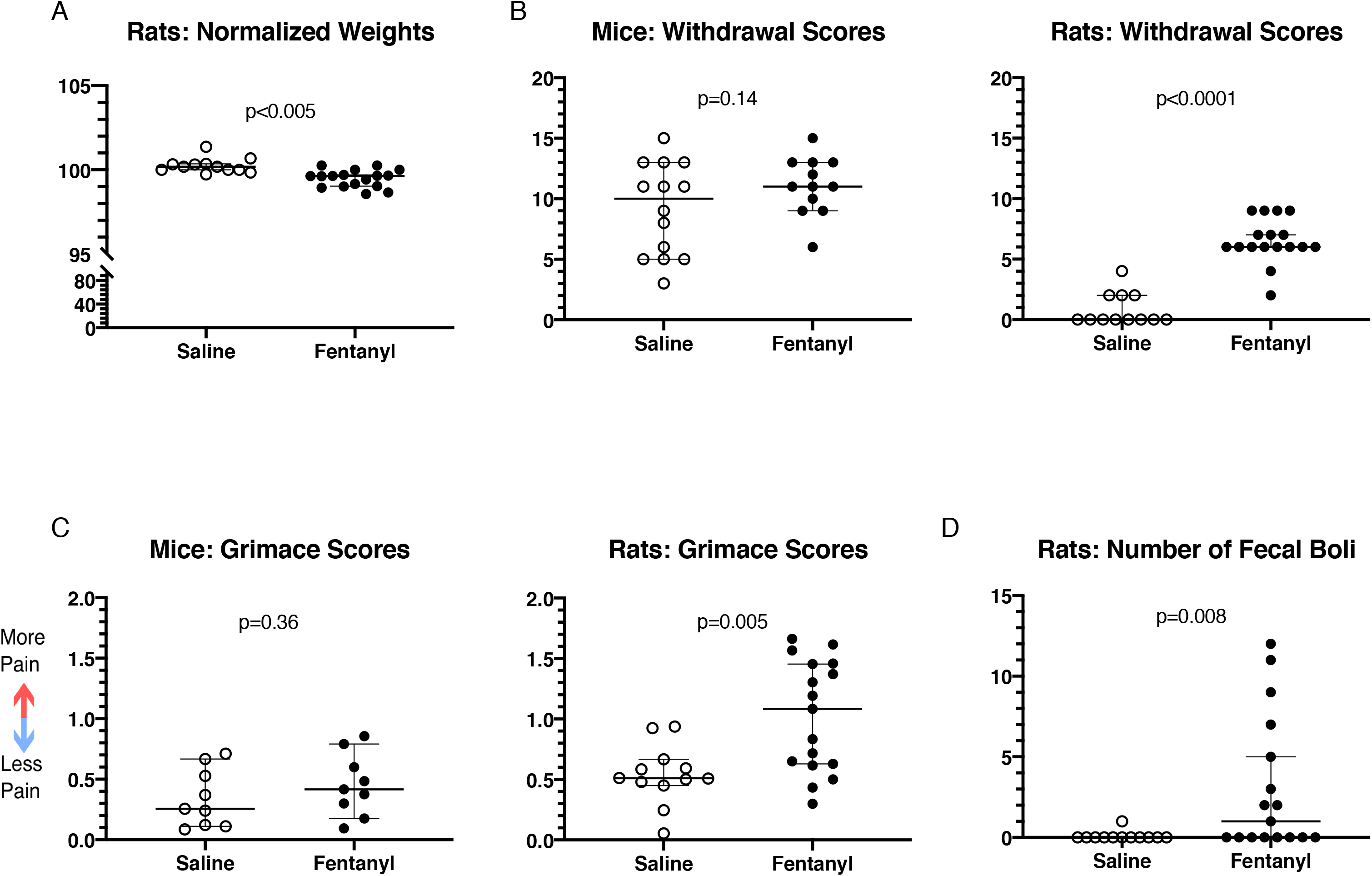
Passive behaviors in mice and rats during precipitated fentanyl withdrawal (A) Rats experience weight loss during precipitated fentanyl withdrawal. (B) Mice and rats both have increased withdrawal scores. (C) Mice experience no change in their grimace scores, while rats display increases. (D) Rats produce more fecal boli during precipitated fentanyl withdrawal. Individual points represent data from a single animal, and horizontal lines depict medians and 95% confidence intervals. Normalized data are expressed as a percent of baseline.

#### Withdrawal Scores

Compared to saline exposed animals, only fentanyl-exposed rats had higher precipitated withdrawal scores. Saline exposed rats (n=12) had median withdrawal scores of 0 (95%C.I. 0 - 2), while fentanyl exposed rats (n=17) had median withdrawal scores of 6 (95% C.I. 6 - 7; Fig. 7B; Cohen’s d = 3.5, Mann-Whitney U = 3, p < 10^−4^). Saline exposed mice (n = 14) had mean withdrawal scores of 9.1 95% C.I. 6.9 - 11.4). Fentanyl exposed mice (n = 12) had mean withdrawal scores of 11(95% C.I. 9.6 - 12.6) (Fig 7B; Cohen’s d = 0.59, t(24) = 1.5, p = 0.14, unpaired t test).

As with spontaneous withdrawal, mice showed multiple somatic withdrawal signs regardless of treatment group (Fig 5B), however more fentanyl treated mice exhibited wet dog shakes after naloxone compared with saline mice (Fig. 5B). In contrast, fentanyl exposed rats showed several withdrawal signs that were largely absent in the saline group (Fig. 5B).

#### Ongoing pain

In fentanyl treated mice, naloxone-precipitated withdrawal did not elicit changes in grimace scores, relative to saline treated controls (Fig. 7C; Saline: n = 9, mean score, 95% C.I. 0.16 - 0.53. Fentanyl: n = 9, mean score 0.45, 95% C.I. 0.26 - 0.65, t(16) = 0.94, p = 0.36, unpaired t test), suggesting no changes in ongoing pain during withdrawal. In fentanyl exposed rats, precipitated withdrawal resulted in higher grimace scores, compared to saline exposed rats (Fig. 7C; Cohen’s d = 1.24, Mann-Whitney U = 40.5, p = 0.005). Saline exposed rats (n = 12) had median grimace scores of 0.51 (95%C.I. 0.45 - 0.67). Fentanyl exposed rats (n = 17) had median grimace scores of 1.1 (95% C.I. 0.63 - 1.5). This suggests that rats in precipitated withdrawal have increased spontaneous pain.

#### Fecal Boli

An often-reported opioid withdrawal sign is increased fecal output, usually diarrhea. Consequently, we examined this metric in rats (platform design precluded this assessment in mice). Fentanyl exposed rats had a higher number of fecal boli during the precipitated withdrawal time period, compared to saline exposed rats (Fig. 7D; Cohen’s d= 0.92, Mann-Whitney U=52.5, p=0.008). Saline exposed rats (n=12) had a median of 0 boli (95% C.I. confined to 0). Fentanyl exposed rats had a median of 1 bolus (95% C.I. 0-5). Notably, all boli were solid and did not qualify as diarrhea.

### Mice show thermal hypersensitivity and rats show increased ongoing pain during precipitated withdrawal

#### Thermal Sensitivity (Heat)

Fentanyl exposed mice had shorter hindpaw withdrawal latencies during precipitated withdrawal, compared to saline exposed animals, suggesting increased sensitivity to heat during withdrawal (Fig 8A; Cohen’s d = 0.64, Mann-Whitney U = 36, p = 0.017. Saline: n = 14, median 107.1%, 95% C.I. 100.6 - 115.4%. Fentanyl: n = 12, median 93.5,, 95% C.I. 87.9 - 104.5%). By contrast, fentanyl exposed rats had higher hindpaw withdrawal latencies during precipitated withdrawal, compared to saline exposed animals, suggesting analgesia to heat stimuli in the fentanyl group (Fig 8A; Cohen’s d=1.06, Mann-Whitney U=14, p=0.0045. Saline: n=8, median 68.7%, 95% C.I. 54.6-121.8%. Fentanyl: n=13, median 127.4%, 95% C.I. 92.8-225.7%).

**Figure 8:**
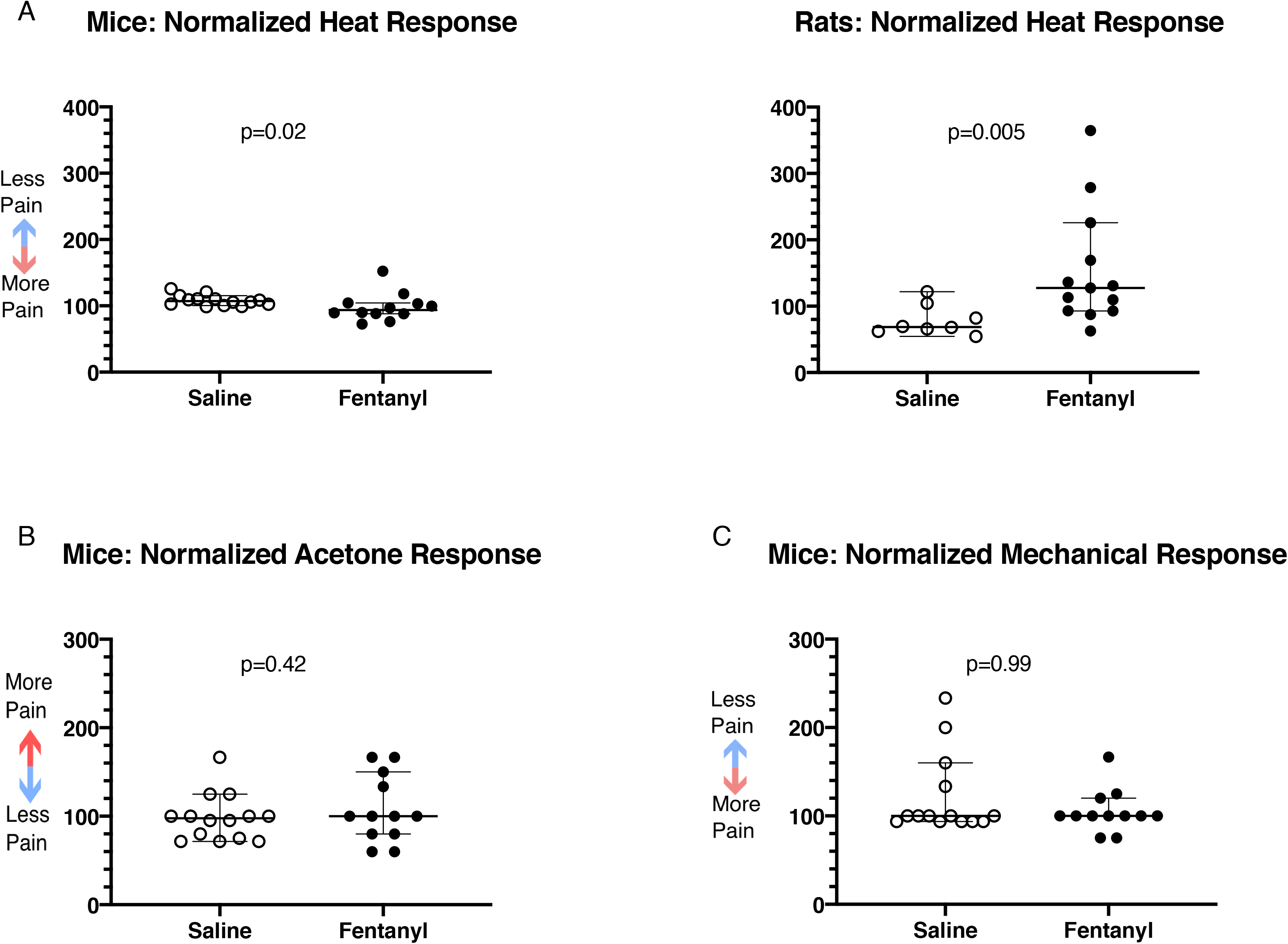
Pain related behaviors during precipitated fentanyl withdrawal. (A) Mice exhibit increased heat sensitivity, while rats have decreased heat sensitivity in precipitated fentanyl withdrawal. (B) Mice experience no change in cold sensitivity. (C) Mice show no change in mechanical sensitivity. Mice experience no change in cold sensitivity. Individual points represent data from a single animal, and horizontal lines depict medians and 95% confidence intervals. Normalized data are expressed as a percent of baseline.

#### Thermal Sensitivity (Cold)

During precipitated withdrawal in mice there was no difference in acetone response scores when comparing saline and fentanyl exposed animals (Fig 8B; Mann-Whitney U = 68, p = 0.42. Saline: n = 14, median 97.6%, 95% C.I. 71.4 - 125%. Fentanyl: n = 12, median 100%, 95% C.I. 80 - 150%). Neither saline nor fentanyl exposed rats responded to acetone application, suggesting that the stimulus was not noxious enough to cause pain in either group.

#### Mechanical Sensitivity

There was no difference in mechanical responses between saline and fentanyl exposed mice (Fig. 8C; Mann-Whitney U=83.5, p=0.99). Both saline exposed mice (n=14) and fentanyl exposed mice (n=12) had median response thresholds that were 100% of baseline values (saline: 95% C.I. 93.8-160%; fentanyl; 95% C.I. 100-120%). The relatively short duration of naloxone, combined with the length of behavioral assessments for the rats, prevented us from recording mechanical sensitivities in this species.

## Discussion

We report somatic withdrawal and pain behaviors in rats and mice after chronic fentanyl exposure under standardized conditions. We report that both rats and mice exhibit increased somatic withdrawal signs during spontaneous and naloxone precipitated withdrawal, although the prevalence of individual behaviors differs between species. The species diverge further in how they express withdrawal-associated pain behaviors: only mice display hyperalgesia evoked by hot and cold stimuli, while rats exhibit increased signs of ongoing, spontaneous pain.

### Intermittent exposure paradigm

Our mice and rats received subcutaneous injections of fentanyl, twice-daily for two weeks, with no exposure on the weekends. We chose the doses of fentanyl in mice and rats based on the LD_50_ for each species, to account for differences in metabolism between the species. We recognize that a full dose-response study is needed to fully account for discrepancies in how each species metabolizes fentanyl.

We used an intermittent delivery schedule to mimic irregular exposure in opioid abusers, where individuals go through repeated bouts of intoxication and withdrawal. Repeated exposure and subsequent withdrawals likely contributed to the outcomes reported here, as they presumably do in human opioid users. Disentangling which symptoms are due to opioid receptor saturation versus the sudden loss of the agonist or the degree to which self-administration contributes to fentanyl withdrawal symptoms, will require additional studies.

### No Sex Differences

We found no sex differences in any fentanyl withdrawal metrics, but recognize that a larger sample size may be necessary to reveal such differences. Indeed, sex differences were reported, in both rats and mice, in both the types and severity of spontaneous and precipitated opioid withdrawal ^47,48^. However, whether sex differences in withdrawal are present depends on the choice of species, behavioral metrics analyzed, and on whether spontaneous or precipitated withdrawal is induced ^49^.

### Mice and rats show somatic signs of withdrawal

In humans, opioid withdrawal is expressed as tachycardia, GI upset, sweating, tremor, restlessness, yawning, anxiety/irritability, pupil dilation, rhinorrhea and lacrimation, bone or joint aches, and piloerection ^2^. While both humans and non-human animals are impacted by affective and emotional components of withdrawal, ^50–53^, here we chose to focus on somatic signs and hyperalgesia. A similar spectrum of withdrawal signs occurs in rat and mouse models of spontaneous and naloxone-precipitated morphine withdrawal ^34,47,54–57^. However, few studies directly compare the withdrawal nuances in different rodent species within the same report. The few studies that do use both species find mostly overlapping withdrawal signs ^58^.

We directly compared fentanyl withdrawal profiles in rats and mice. While we find some overlap of signs in our model of fentanyl withdrawal, their prevalence varied by species. For mice in both spontaneous and precipitated withdrawal, grooming, wet dog shakes, teeth chattering, swallowing, genital-specific grooming, and paw tremors were the most prevalent behaviors in the fentanyl group (Fig. 5). For these mice, a higher frequency of wet dog shakes appears to be the behavior that best indicates withdrawal from the intermittent fentanyl exposure paradigm. In comparison, rats showed considerably fewer withdrawal-associated behaviors at baseline: Only wet dog shakes and hyperirritability were present in more than one rat at baseline. In rats in spontaneous withdrawal, abdominal constrictions or writhing, wet dog shakes, teeth chattering, and swallowing were the most prevalent behaviors in fentanyl treated animals (Fig. 5). During precipitated withdrawal in rats, we observed animals in plexiglass chambers, due to the limited time in which to complete all tests and observations before naloxone’s effects waned. Therefore, wet dog shakes and abdominal constrictions or writhing were limited by space constraints. In these conditions, teeth chattering, swallowing, ptosis, and hyperirritability were the most prevalent behaviors in fentanyl treated rats, distinguishing them from their saline treated counterparts.

Although jumping behavior in mice, and weight loss in both species are considered highly reliable opioid withdrawal metrics ^20,56,59^, the design of our observation chambers—specifically, their short height—limited jumping behavior expression in our study, potentially accounting for the discrepancy between the literature and our results.

### Mice and rats show weight loss during chronic opioid administration and spontaneous withdrawal

During spontaneous withdrawal, fentanyl exposed rats and mice both lost weight. This may be an effect of withdrawal, or of repeated drug exposure, since chronic fentanyl and morphine exposure (both intermittent and continuous) lead to weight loss ^32,33,60,61^.

### Rats show weight loss and moderately increased fecal boli production during precipitated withdrawal

Fentanyl exposed rats lost weight over the course of precipitated withdrawal (approximately 1 hour maximum), presumably because they produced more fecal boli than saline animals during this period. Notably, these boli did not constitute profuse diarrhea, as reported in studies using different opioids and exposure routes ^55,62,63^.

### Rats show signs of ongoing pain during withdrawal

Humans experience stimulus-evoked withdrawal hyperalgesia, but the most common descriptions of opioid withdrawal pain includes “muscle and bone aches” ^2^. In contrast, data assessing ongoing, spontaneous pain in rodents during opioid withdrawal is lacking. Here, we used a well-validated, non-reflexive measure of spontaneous pain in rodents, the grimace scale ^40–42^ to assess this quality of pain in animals undergoing fentanyl withdrawal. Fentanyl exposed rats have higher grimace scores during both spontaneous and precipitated withdrawal, suggesting that they experience more ongoing pain. Mice showed no differences in grimace scores during withdrawal, suggesting that they are either free of spontaneous pain during fentanyl withdrawal, or, due to social dynamics, are less likely to display visible signs of pain or distress.

### Mice and rats show no signs of mechanical hyperalgesia during fentanyl withdrawal

Mechanical hyperalgesia following opioid exposure in rodents has been observed using the paw pressure vocalization test ^24,64–68^, tail pressure test ^69^, or with von Frey testing ^70^. Using 10-trial von Frey testing for rats, and a 3-trial testing paradigm for mice (see Methods), we did not observe any withdrawal-induced changes in mechanical sensitivity. Previous studies ^24,64,66,70^ were designed to model *acute* fentanyl exposure, administering lower doses of fentanyl during one exposure day. These studies yielded hyperalgesia emerging several hours after opioid-induced analgesia had waned, and lasting several days post-exposure. In contrast, our study involves fentanyl injections that escalated to higher total dosages, are more dispersed over time, and continue across 2 weeks. The absence of mechanical hyperalgesia at 20-22 hours spontaneous withdrawal in our study may reflect differences in our animal subjects or exposure paradigm, which was intended to model patterns of fentanyl misuse and abuse instead of acute exposure.

### Mice show thermal (heat) hyperalgesia during fentanyl withdrawal

Previous studies demonstrated hyperalgesia to heat stimuli following opioid exposure using the tail flick ^18–20,66^, hot plate ^21,22,71–73^ or Hargreaves tests ^18,23,69^. Consistent with prior reports ^73^, fentanyl exposed mice developed thermal hyperalgesia at both 20-22 hours spontaneous withdrawal and during precipitated withdrawal. In our hands, rats showed no signs of heat hyperalgesia. In fact, during precipitated withdrawal, fentanyl exposed rats had longer withdrawal latencies compared to saline animals. As we collected heat responses last during precipitated withdrawal experiments in rats, it is possible that, by this time (30-60 minutes after naloxone), precipitated withdrawal hyperalgesia may have faded or been overshadowed by returning fentanyl analgesia, since naloxone was administered 5 minutes after fentanyl. The species difference might also reflect the different thermal assays we used for rats and mice. The Hargreaves test which we used in rats is thought to primarily reflect spinal mechanisms, whereas the hotplate test which we used in mice also reflects activity in supraspinal pathways ^74^.

### Mice show thermal (cold) hyperalgesia during fentanyl withdrawal

In humans, cold hyperalgesia occurs during chronic opioid use and withdrawal ^11,75,76^. In rodents, evidence of cold hyperalgesia during withdrawal is less extensive. Thus, our effort to assess this metric adds important data to the literature on altered pain perception during fentanyl withdrawal. Morphine withdrawal hyperalgesia has been shown in male C57Bl6/J mice tested on a cold plate ^77^. One group reported minimal effects of chronic fentanyl exposure and withdrawal on cold sensitivity in aged rats ^60^. Here, we use the acetone test to measure cold sensitivity ^78,79^. In our study, only fentanyl exposed mice in spontaneous withdrawal showed cold hyperalgesia. Surprisingly, mice undergoing precipitated withdrawal did not. No rats, including the saline group, responded to acetone application in our study. Compared to other investigations, our rats are older, with presumably thicker plantar skin. Whether this physical difference impaired the cold response, or whether there is no cold hyperalgesia associated with this model of withdrawal remains to be determined.

### Limitations

Our study seeks to understand how mice and rats model signs of withdrawal from a repeated, intermittent fentanyl exposure paradigm that mimics patterns of human fentanyl misuse, and to compare how pain-associated metrics are altered in these conditions. While we find distinct profiles of withdrawal behaviors and associated pain in mice compared to rats, we note that there are methodological differences that can contribute to these outcomes. We based our fentanyl dosing regimen on the LD50s for each species, resulting in mice receiving a higher dose than rats. A full dose-response experiment for each species would provide a more complete picture of how this difference impacts withdrawal signs in mice and rats. Further, the different methods used for each species to test for changes in mechanical and thermal sensitivities capture different aspects of these modalities. Thus, while our data suggest that mice and rats show different withdrawal signs and pain-associated behaviors during spontaneous and precipitated withdrawal from intermittent fentanyl exposure, the experimental factors outlined above are important to consider when interpreting these results.

### Conclusion

Using escalating, intermittent, subcutaneous fentanyl exposure to evaluate withdrawal and associated pain in mice and rats, we find only partially overlapping somatic withdrawal signs in both species and thermal withdrawal hyperalgesia in mice. Rats show signs of ongoing pain during withdrawal while these signs were not observed in mice. These data support the use of rodents in modeling aspects of opioid withdrawal and highlight that species choice is important when designing experiments to further our understanding of opioid withdrawal and withdrawal-associated pain.

## Authors Contributions

NC, AK, and OU were responsible for study design and concept. KA, NC, MEF, CJ, and OU collected and analyzed data. NC and OU drafted the manuscript with critical revisions and input from MEF and CJ. All authors reviewed content and approved the final version.

## Acknowledgments

We are thankful to Andrew Furman for sharing his MATLAB programs for Grimace Analysis. This study was funded by the Opioid Use Disorders initiative of MPowering The State (State of Maryland) to AK, and by the National Institutes of Health K99DA050575 to MEF. The content is solely the responsibility of the authors and does not necessarily represent the official views of the National Institutes of Health. The funding sources had no role in study design; the collection, analysis and interpretation of data; the writing of the report; or in the decision to submit the article for publication.

